# Neural defensive circuits underlie helping under threat in humans

**DOI:** 10.1101/2021.12.17.473167

**Authors:** Joana B. Vieira, Andreas Olsson

## Abstract

Empathy for others’ distress has long been considered the driving force of helping. However, when deciding to help others in danger, one must consider not only their distress, but also the risk to oneself. Whereas the role of self-defence in helping has been overlooked in human research, studies in other animals indicate defensive responses are necessary for the protection of conspecifics. In this pre-registered study (N=49), we demonstrate that human defensive neural circuits are implicated in helping others under threat. Participants underwent fMRI scanning while deciding whether to help another participant avoid aversive electrical shocks, at the risk of also being shocked. We found that higher engagement of neural circuits that coordinate fast escape from self-directed danger (including the insula, PAG and ACC) facilitated decisions to help others. Importantly, using Representational Similarity Analysis, we found that the strength with which the amygdala and insula uniquely represented the threat to oneself (and not the other’s distress) predicted helping. Our findings indicate that in humans, as other mammals, defensive mechanisms play a greater role in helping behaviour than previously understood.

## Introduction

Helping someone in danger (for example, by saving a person who fell on the train tracks, or running into a building in flames to rescue someone inside) may expose oneself to health and life-threatening risks. Nevertheless, such helping behaviors are observed across species (*1–5*). Risky helping differs from other altruistic actions in that it occurs in the simultaneous presence of two highly salient cues: the distress of a conspecific in need, and a potential threat to the self. In humans, a wealth of research has been dedicated to the former, explaining how perceiving distress in others may trigger the motivation to help (*6, 7*), particularly if the helper is not under threat themselves (*3*). But virtually nothing is known about how, in a threatening situation, one’s own responses to the threat may drive decisions to help. More so, animal research suggests defensive brain mechanisms may in fact be implicated in aiding or protecting conspecifics (*8–10*). Understanding the neurocognitive processes underlying the motivation to both safeguard oneself and help others is critical to explain inter-individual behaviour in dangerous contexts. The overarching goal of our study was thus to determine how one’s own defensive responses to threat guide decisions to help others in dangerous situations.

In humans, defensive responses to threat are graded as a function of the proximity or imminence of the threatening stimulus, paralleling predatory avoidance responses in other mammals (*11, 12*). Distal and unpredictable threats are typically associated with risk assessment and intermittent anxiety, allowing for slower and more flexible escape decisions. As threat imminence increases and an attack becomes more likely, fixed and species-specific responses are triggered, such as such as freezing or, if immediate avoidance is necessary, fight-or-flight. Some behavioural reports indicate that different states along the defensive continuum may have dissociable effects on prosocial behaviour. For example, following acute social stress, participants behave more prosocially in economic games (*13*–*15*), make more moral decisions (*16*), and show greater empathy for others (*13*). Importantly, it has been shown that individuals were more likely to help a co-participant avoid aversive electrical shocks when the threat of shock was imminent rather than distal (*17*). This behavioural pattern was accompanied by faster reaction times and heart rate during imminent compared to distal threats, paralleling what has been found in response to imminent self-directed threats (*18, 19*). Consistent with these laboratorial studies, higher danger in real life situations (captured via public surveillance footage) has been associated with higher likelihood of bystander intervention (*20*). Taken together, these findings suggest that defensive states triggered by high threat imminence may not only enable fast avoidance of self-directed threats, but also motivate helping when others are under threat. Yet, the neural basis of these effects are unclear. Specifically, it is unknown how the activation of specific sub-circuits underlying different defensive states (e.g., freezing versus fight-or-flight) impacts decisions towards others in a threatening context.

We aimed to characterize the involvement of different defensive neural responses on helping under threat. To do so, we used a paradigm adapted from Vieira et al. (2020), in which participants make helping decisions at different stages of threat imminence (details in Figure 1A). Briefly, a participant is asked to decide whether or not to help a co-participant (in reality, a confederate) avoid aversive electrical shocks. In each trial, the participant watches a supposedly live video-feed of the co-participant, and a visual cue signaling an upcoming shock. The participant is asked to decide whether to help the co-participant avoid the shock at the risk of receiving a shock her/himself. These decisions are prompted in some trials in the beginning of the trial (*distal threat),* and in others immediately prior to the shock delivery (*imminent threat*). If the participant decides not to help, the co-participant always receives a shock; if the participant decides to help, there is a fixed probability both participant and co-participant will receive a shock.

**Figure 1.**
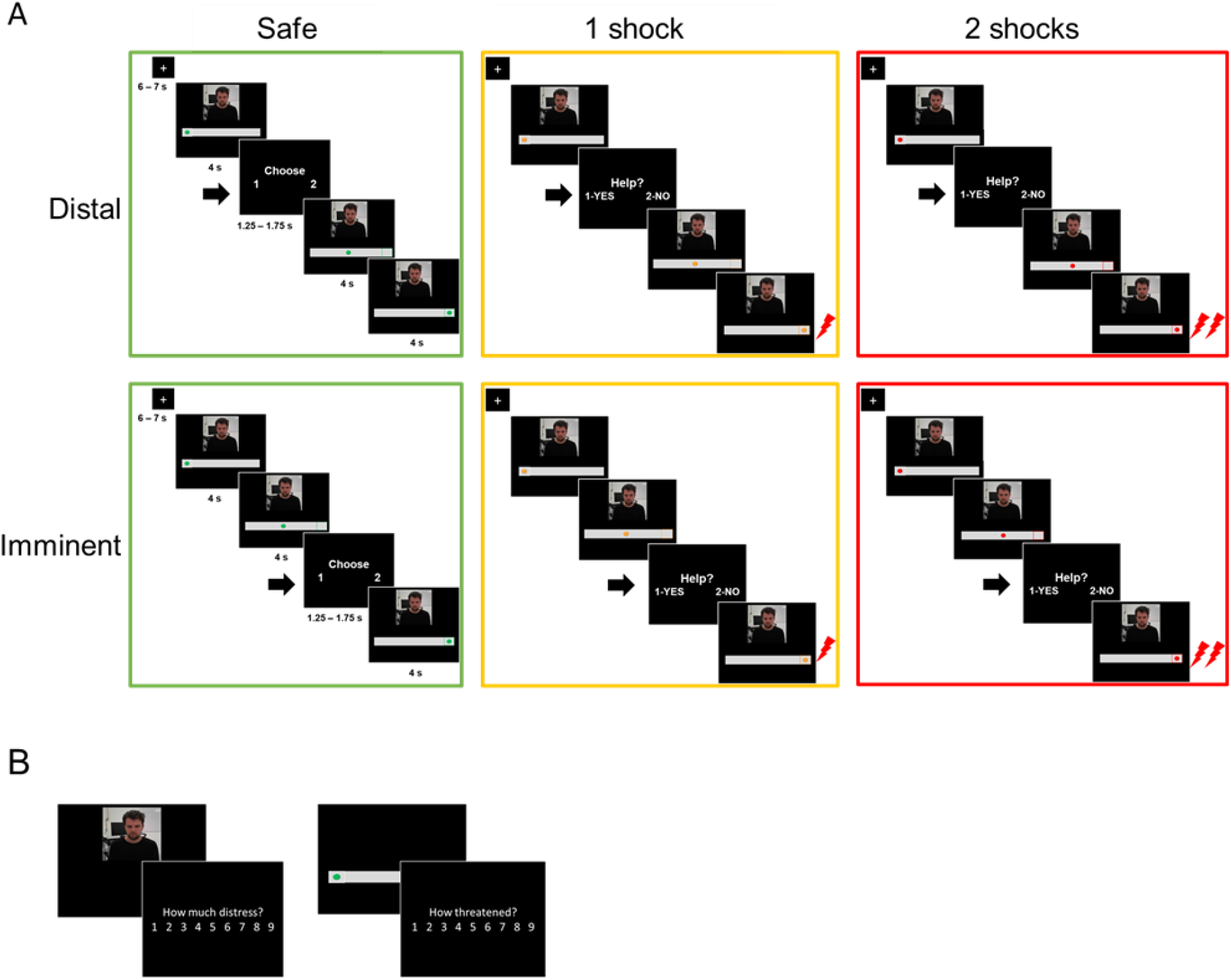
Outline of the experimental tasks. A. fMRI helping under threat task. Participants saw the co-participant on the screen, together with a visual cue signaling threat (an upcoming shock). There were three threat levels: safe (0 shocks, green circle), moderate threat (1 shock, yellow circle) and high threat (2 shocks, red circle). In each trial of the task, the circle started static on the left (4 s), and then moved to the right (4 s). Participants were prompted to decide whether they wanted to help the co-participant or not (1.25 – 1.75 s) either in the beginning of trial (distal) or right before the moment of shock delivery (imminent). Therefore, the available time to make a decision was identical in distal and imminent threats. If participants decided to help, there was a 70% chance both themselves and co-participant would receive shocks; if they decided not to help, the co-participant would always receive a shock, and the participant would not. Decisions prompted on safe trials were to arbitrarily choose to press 1 or 2, since no shocks would be administered. B. After the fMRI task, outside the scanner, participants re-watched clips of the co-participant presented during the scan, and were asked to rate how much “discomfort, anxiety or uneasiness” he was experiencing in each clip on a 9-point scale. They were also presented images of the threat cues and asked to rate, on the same scale, how threatened they felt themselves when they saw those images during the scan.

Our hypotheses were guided by previous work on neural responses to the imminence of self-directed threats. It has been shown that the response to distal threats (*i.e.*, unpredictable, spatially distant, retreating, slow moving) is coordinated by so-called “cognitive fear” circuits, which include the ventromedial prefrontal cortex (vmPFC) and hippocampus (*12, 21*). Conversely, imminent threats (*i.e.,* predictable, spatially close, looming, fast moving) predominantly engage “reactive” fear circuits, which include the dorsal anterior cingulate (dACC), insula, and periaqueductal gray (PAG) (*12, 21*). The amygdala plays a central role in both circuits, namely by coordinating adaptive switches between defensive states as a function of threat imminence (for example, from freezing to fight-or-flight), through oxytocin-mediated communication between its central (CeA) and basolateral nucleus (BLA) (*22, 23*). Based on these findings, we expected neural activation within the full defensive circuitry to respond to the threat level of the trial (i.e., safe, 1 shock and 2 shocks), with higher engagement of brain regions previously included in cognitive fear circuits (i.e., vmPFC and hippocampus) in response to distal threats, and higher engagement of regions included in reactive fear circuits (i.e., insula, dACC and PAG) in response to imminent threats. At the behavioural level, prior findings (*17*) showed that helping decisions were more frequent under imminent than distal trials, suggesting that the activation of reactive fear circuits would facilitate helping behaviour. We thus predicted that helping decisions would be associated with higher engagement of brain regions included in reactive fear circuits (*i.e.* amygdala, insula, ACC and PAG).

One important aspect of our paradigm was that, as in most real-life dangerous situations, the threat and the conspecific in need were simultaneously presented. To dissociate representations of threat and other’s distress within the defensive circuitry, and assess their unique role on helping behaviour, after the scan we asked participants to rate the degree of distress experienced by *the co-participant* in each clip showed during the scan; also, participants rated how threatened they felt *themselves* when they saw the visual threat cues during the scan. These ratings were used as behavioural models in a representational similarity analysis (RSA; see Materials and methods) that identified neural representations of other’s distress and of threat to the self, and determined their association with helping behaviour.

The demonstration that neural representations of other’s distress are positively associated with helping decisions would support existing empathy-based explanations of helping. Indeed, it has been proposed that helping a conspecific in danger results primarily from an evolutionarily preserved motivation to care for offspring in mammals, which is triggered by signals of distress and vulnerability, and is especially likely to occur *if* the helper is not under threat themselves (*3*). However, evidence in rodents indicates that, rather than conflicting, defensive responses may be *required* for helping and caregiving: e.g., anxious rat mothers display enhanced maternal behavior after pharmacological activation of defensive brain circuits (*9, 24, 25*), whereas mice bred to have low anxiety display significant defects in maternal behaviors (*26*); and helping behavior in rats is compromised following treatment with anxiolytic drugs that suppress defensive circuits (*8*). According to these animal findings, an alternative prediction to empathy-based accounts is that the neural representation of threat to the self would promote helping of others.

## Results

### Helping decisions did not vary based on threat imminence or threat level

Using Generalized Linear Mixed Models (GLMMs), we found no significant effect of either threat imminence (*β*=.017, *se*=.014, *t*=1.24, *p*=.218), threat level (*β*=-.068, *se*=.053, *t*=-1.282, *p*=.205), nor a threat imminence*level interaction (*β*=-.099, *se*=.019, *t*=-.497, *p*=.622) on the percentage of helping decisions throughout the task (Figure 2A). Despite the lack of group-level effects of threat imminence (which were predicted based on previous work;, *17*), individual data showed that the number of participants helping more during imminent than distal threats was higher (n=27) than those helping more during distal (n=11), or the same amount during imminent and distal (n= 11) (Figure 2B).

**Figure 2.**
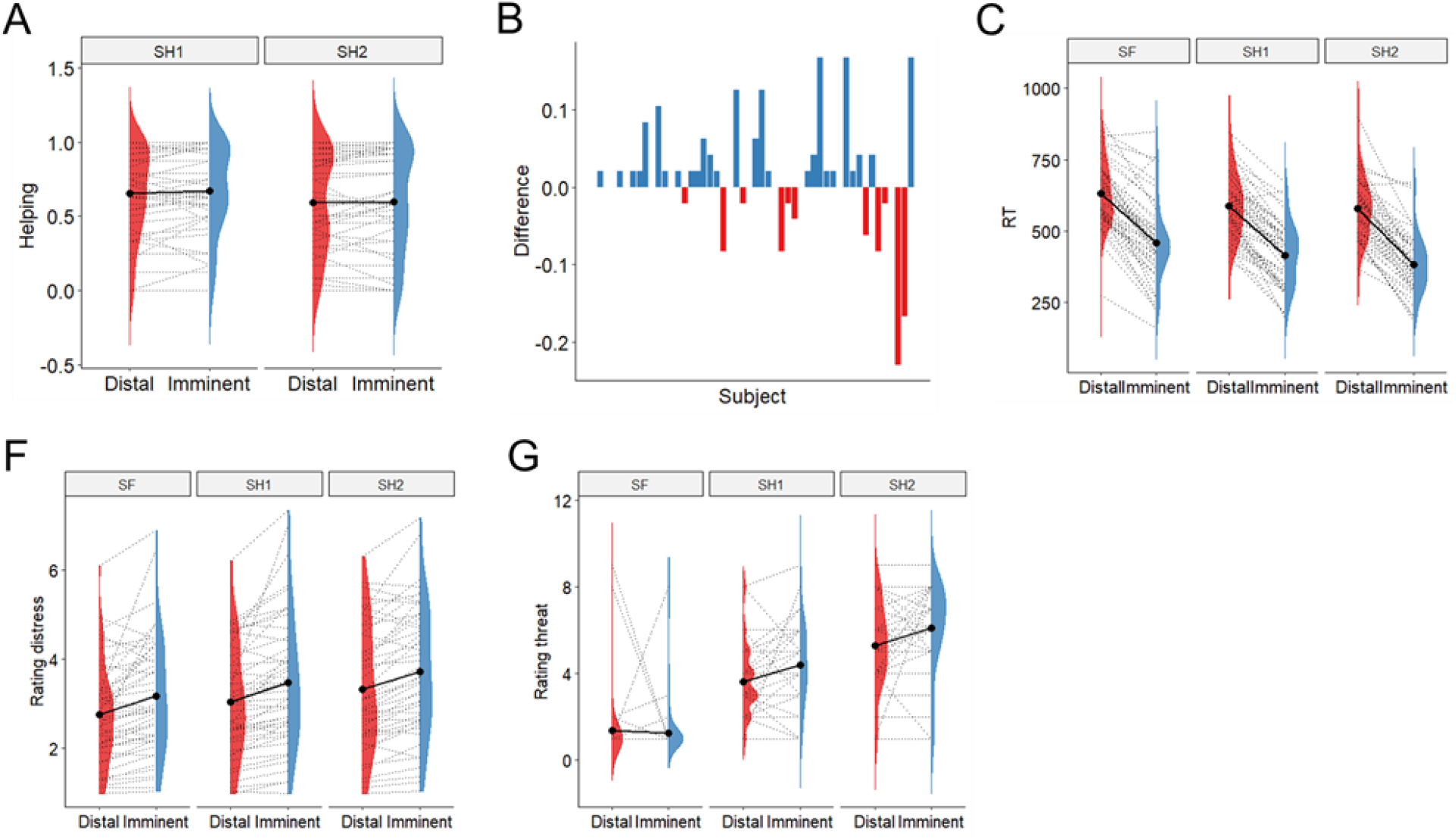
A. There was no evidence of differential helping during imminent and distal threats, nor during 1-shock and 2-shock trials. B. Difference between proportion of helping in imminent and distal trials (y axis) across subjects (x axis); 27 participants helped more during imminent than distal threats, 11 helped more during distal, and 11 helped the same amount. C. Responses were faster during imminent than distal trials across threat levels. D. Participants rated the co-participant’s distress as higher during imminent than distal trials, and as progressively higher across the three threat levels. E. Participants reported feeling more threatened when watching 2-shock cues (red circle), followed by 1-shock cues (yellow circle) and safe cues (green circle), and when watching cues signaling imminent than distal threat.

Previous work also suggested the impact of threat imminence on helping might vary based on empathic tendencies (*17*). We thus checked whether there was an interaction between the empathic concern scale of the Interpersonal Reactivity Inventory (*27*) and threat imminence on helping. When empathic concern was added in the model, results showed no significant effects of threat imminence (*β*=.018., *se*=.014, *t*=1.249, *p*=.215) and threat level (*β*=-.050, *se*=.051, *t*=-.985, *p*=.330), and no significant interaction between imminence and empathic concern (*β*=.018, *se*=.014, *t*=1.292, *p*=.20). However, a significant association emerged between empathic concern and helping behaviour (*β*=-.101, *se*=.041, *t=-* 2.473, *p*=.02), indicating those higher in empathic concern displayed less frequent helping behaviour.

Also, to account for the possibility that decisions varied throughout the experiment (e.g., participants helped more in the beginning than towards the end), we also performed a mixed effects logistic regression on single trial dichotomous responses (help or no help), including the trial number as a fixed effect. This analysis revealed no significant effects, indicating the individuals did not respond differently as time passed.

Finally, in line with previous work (*17*), analysis of reaction times showed individuals made faster decisions during imminent versus distal trials (*β*=-173.30, *se*=12.09, *t*=-14.33, *p*<.0001), and for shock versus safe trials (*β*[1 shock]=-43.26, *se*=12.76, *t*=-3.39, *p*=.001; *β[2* shock]=-53.35, *se*=13.43, *t*=-3.97, *p*=.0002), with no significant threat imminence*level interaction (Figure 2C).

### Participants were sensitive to variations in the co-participant’s distress, threat imminence, and threat level (manipulation check)

After the scan, participants were asked to re-watch all clips of the co-participant during the scan, and rate the level of “discomfort, anxiety or uneasiness” they thought *he* was experiencing in each clip. Of note, these clips were shown without the threat cues (see Figure 1B) that were also present during the scan, in order to isolate the response to distress and threat. Results showed participants rated the distress of the co-participant being significantly higher during imminent than distal clips (*β*=.418, *se*=.089, *t*=4.676, *p*<.0001), and progressively higher across the three levels of threat (1 Sh: *β*=.279, *se*=.077, *t*=3.611, *p*=.0005; 2 Sh: *β*=.559, *se*=.081, *t*=6.853, *p*<.0001; reference class was safe). No significant threat level*imminence interaction was found (Figure 2F). These results suggest the video clips used in the scan successfully portrayed subtle variations in cues of distress by the co-participant.

Participants were also presented isolated images of the threat cues used during the scanning task (namely, the green, yellow and red circles, both in the distal and imminent positions; Figure 1B), and asked to rate how threatened they felt *themselves* when they saw those visual cues during the scan. Results showed participants rated threat stimuli as more threatening (1 Sh: *β*=2.255, *se*=.297, *t*=7.598, *p*<.0001; 2 Sh: *β*=3.92, *se*=.297, *t*=13.89, *p*<.0001; reference class was safe; Figure 2G). Additionally, imminent cues were rated as more threatening, but only for 1 and 2 shocks, and not for safe trials (imminence*1 Sh: *β*=.894, *se*=.420, *t*=2.129, *p*=.034.; imminence*2 Sh: *β*=.936, *se*=.420, *t*=2.23, *p*=.027).

### Neural responses

Analysis of brain responses focused on a liberally defined set of brain regions that integrate the brain’s defensive system, namely the vmPFC and vlPFC/IFG (*28*–*30*), the hippocampus (*21*), the insula (*29, 30*), the ACC (*28, 29*), the amygdala (*22, 28*) and the midbrain (*21, 28–31*) (Figure 3).

**Figure 3.**
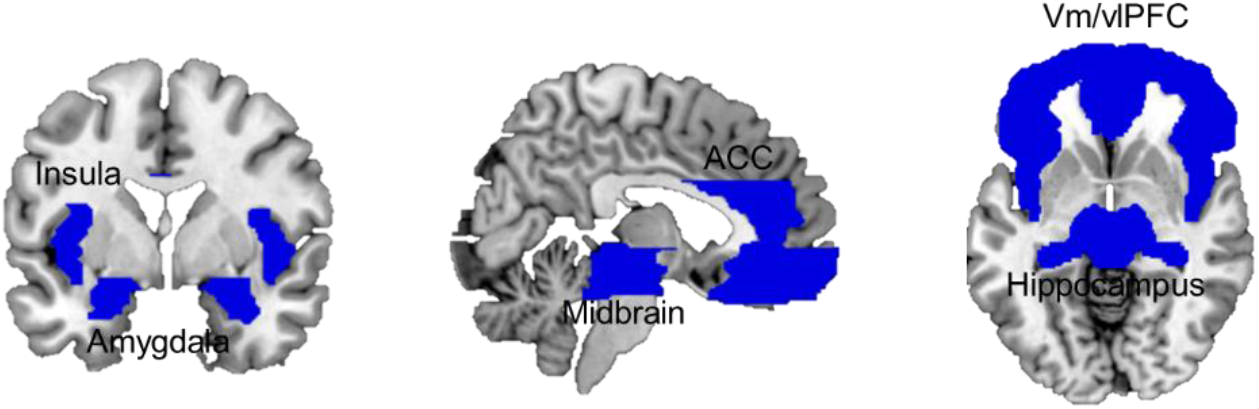
Combined ROI mask including bilateral ventral and lateral medial frontal cortex, dorsal ACC, insula, hippocampus, amygdala and midbrain.

#### a. Multivariate and univariate differentiation of threat imminence and level in the defensive circuitry

We performed a Support Vector Machine regression to identify sites in which activation patterns were linearly associated with increasing threat level (*i.e.,* from safe, to 1 shock and 2 shocks). As predicted, results showed that throughout the defensive circuitry (i.e., amygdala, insula, ACC, hipoccampus, and regions within the orbitofrontal cortex) multivariate activation tracked with threat level (*FWE* < .05, *k* > 10; Figure 4B).

**Figure 4.**
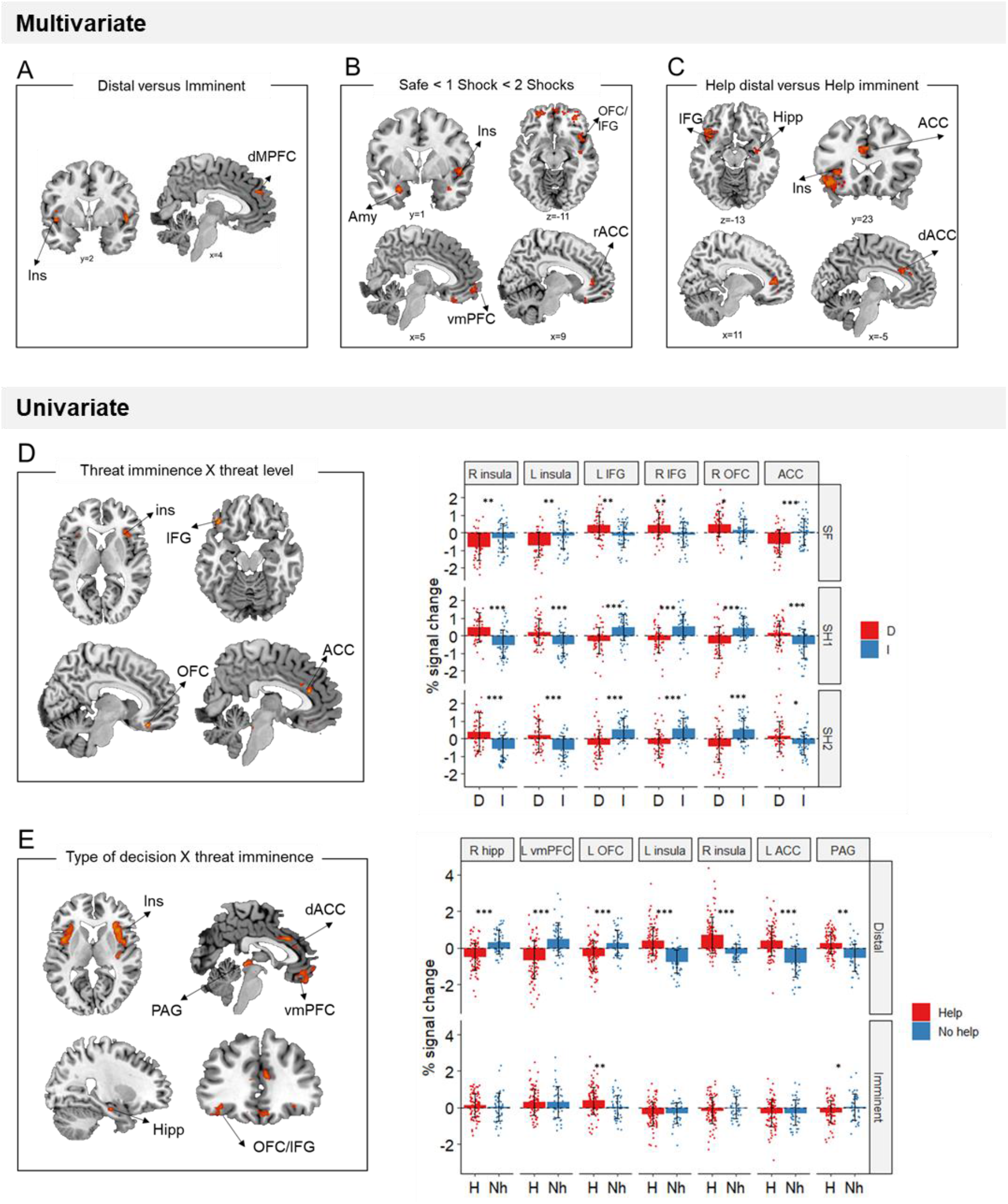
A. Local multivoxel activation patterns (identified by searchlight analysis) in the insula and dmPFC were distinguishable between distal and imminent threats, irrespective of helping decisions. B. Local multivoxel activation patterns (identified by support vector regression) in the amygdala, insula, OFC/IFG, vmPFC, and ACC were linearly associated with varying threat level. C. Local multivoxel activation patterns (identified by searchlight) in the insula, IFG, hippocampus and ACC were distinguishable when making helping decisions under distal and imminent threat. D. Clusters in the insula, IFG, OFC and ACC displayed a significant threat imminence*threat level interaction. E. Clusters in the insula, ACC, IFG/OFC, vmPFC, hippocampus and PAG displayed a significant type of decision*threat imminence interaction. Ins: insula; ACC: anterior cingulate cortex; AMY: amygdala; OFC: orbitofrontal cortex; IFG: inferior frontal gyrus; vmPFC: ventromedial prefrontal cortex; PAG: periaqueductal gray; Hipp: hippocampus; dmPFC: dorsomedial prefrontal cortex. \**p*<.05, \*\**p*<.01, \*\*\**p*<.001.

Additionally, to identify brain regions with differential average activation to the imminence of threatening stimuli specifically, we ran a univariate threat imminence (distal, imminent) by threat level (safe, 1 shock, 2 shocks) ANOVA. Brain regions displaying a significant threat imminence by threat level interaction included the bilateral insula, OFC and IFG and ACC (Table 1; Figure 4D). In shock trials (both 1 and 2 shocks), the bilateral insula and ACC presented higher activation for distal compared to imminent threats, whereas the bilateral IFG and right OFC showed higher activation in imminent compared to distal threats (full ANOVA results in Table S1). These results were opposite to our predictions that brain regions previously implicated in reactive fear circuits (*i.e.,* insula and ACC) would be more active during imminent threats, and regions implicated in cognitive fear circuits (*i.e.,* IFG and OFC) during distal threats. Finally, using a Searchlight cross-classification algorithm (12mm-radius sphere), we also identified brain sites in which multivariate patterns were distinguishable between distal and imminent threats. We found that multivariate patterns in the bilateral insula and dorsal medial prefrontal gyrus dissociated between distal and imminent threats (*FWE* < .05, *k* > 10; Figure 4A).

**Table 1.** Multivariate results based on threat imminence (distal – imminent) and level (safe, 1 shock, 2 shocks; FWE < .05).

| **Searchlight Distal versus Imminent** |  |  |  |  |  |
| --- | --- | --- | --- | --- | --- |
|  | **R/**<br>**L** | **k** | **x, y, z** | **T** | **BA** |
| Insula, superior temporal gyrus | R | 40 | 46, -4, -8 | 7.43 | 22 |
| Insula | L | 17 | -46, 2, -4 | 6.64 | 13 |
|  |  | 18 | -42, 4, 8 | 7.15 |  |
| Medial prefrontal cortex | R | 20 | 10, 48, 28 | 7.13 | 9 |
| **SVM regression SF - 1 SH - 2SH** |  |  |  |  |  |
| Hippocampus | R | 12 | 24, -10, -20 | 7.56 |  |
| Insula | R | 25 | 42, -8, -8 | 8.34 | 13 |
| Rolandic operculum | R | 139 | 52, 0, 0 | 7.12 | 22, 47 |
| Superior temporal gyrus, amygdala | R | 11 | 38, 2, -24 | 7.73 |  |
| Amygdala | L | 37 | -24, 4, -24 | 7.96 |  |
| Rectus | R | 12 | 6, 30, -24 | 7.07 | 11 |
| Anterior cingulate | R | 13 | 8, 38, 8 | 7.02 |  |
| Middle orbital frontal gyrus | R | 42 | 32, 44, -14 | 7.32 | 11 |
| Middle orbital frontal gyrus | L | 11 | -4, 48, -10 | 6.75 | 11 |
|  |  | 44 | -28, 54, -10 | 7.59 |  |
|  |  | 41 | 40, 56, -4 | 7.18 |  |
| Superior frontal orbital gyrus | R | 19 | 16, 54, -14 | 6.78 | 10 |
| Medial frontal gyrus | R | 30 | 6, 58, -8 | 6.97 | 10 |
| **Searchlight Help during distal versus imminent threats** |  |  |  |  |  |
|  | R | 29 | 36, -8, -16 | 8.04 |  |
| Hippocampus | R | 21 | 50, 4, -2 | 7.01 | 22 |
| Insula | R | 23 | 32, 14, 14 | 7.18 | 13 |
| Inferior frontal gyrus | L | 207 | -48, 20, -6 | 8.81 | 47, 38, 22, 13, 45 |
| Mid cingulate, dorsal anterior cingulate | R | 43 | 2, 22, 30 | 7.25 | 32, 9, 24, 6 |
| Insula | L | 45 | -32, 26, 0 | 7.83 | 47, 13, 45 |
| Anterior cingulate | R | 53 | 10, 40, 8 | 7.92 | 32, 10 |

#### b. Greater engagement of reactive fear circuits led to helping

To test our prediction that higher engagement of reactive fear circuits would lead to helping, we performed a decision type (help, not help) by threat imminence (distal, imminent) ANOVA, and focused on brain regions displaying a significant interaction between the two (which would indicate activation differences when making decisions under distal and imminent threat). Of note, due to the reduced number of not helping trials, ‘no help’ decisions in this analysis included not only threat trials in which participants did not help, but also responses made in safe trials (details in the Materials and methods). Results showed a significant interaction in the midbrain PAG, bilateral insula, right hippocampus, dorsal ACC, OFC and vmPFC, which was driven by the distal condition (Table, 2; Figure 4E). Indeed, during distal threats, higher activation in the hippocampus, vmPFC and OFC was followed by decisions not to help, whereas higher activation in the dACC, PAG and insula led to helping decisions. We additionally ran a Searchlight analysis to localize dissociable neural patterns guiding decisions under distal and imminent threats. Results showed that, prior to helping decisions, patterns of activation in the insula, hippocampus and dorsal cingulate were distinguishable between distal and imminent threats (*FWE* <.05, *k* > 10; Figure 4C).

**Table 2.** Results of the univariate ANOVAS (FWE <.05).

| Univariate threat level*imminence interaction |  |  |  |  |  |
| --- | --- | --- | --- | --- | --- |
|  | R/L | k | x, y, z | F | BA |
| Insula | R | 768 | 38, 20, 6 | 28.14 | 13 |
| Insula | L | 521 | -30, 24, -4 | 20.58 | 13 |
| IFG, OFC | L | 593 | -40, 32, -16 | 23.72 | 11 |
| OFC | R | 407 | 8, 32, -20 | 23.43 | 11 |
| ACC | R | 278 | 4, 32, 20 | 18.29 | 24 |
| OFC, IFG | R | 52 | 34, 40, -12 | 13.18 | 11, 47 |
| Type of decision*imminence interaction |  |  |  |  |  |
|  | R/L | k | x, y, z | F | BA |
| Midbrain |  | 60 | 2, -32, -4 | 16.85 |  |
| Insula | R | 192 | 38, -16, -2 | 20.62 | 13, 47, 22, 44, 6, 45, 21 |
| Hippocampus | R | 38 | 30, -14, -20 | 15.03 |  |
| Insula | L | 595 | -36, -8, -4 | 18.68 | 13, 22, 44, 6, 47, 45 |
| Dorsal anterior cingulate | L | 151 | -2, 14, 30 | 16.13 | 24, 6, 32, 5, 4 |
| Insula | R | 748 | 30, 20, -8 | 18.18 | 47, 13, 22, 44, 6, 45, 21 |
| Inferior frontal/orbital gyrus | L | 46 | -34, 36, -10 | 11.47 | 11 |
| Rectus, ventral medial frontal gyrus | L | 238 | 0, 46, -20 | 19.94 | 11, 10, 25 |

#### c. Neural representations of threat promoted helping

One of our goals was to determine whether helping decisions were predominantly driven by the response to another person’s distress, by one’s own defensive state, or by both. To this end, we performed Representational Similarity Analysis (RSA; *32*). This analysis was done separately for imminent and distal trials, and comprised 3 steps (see Figure 5 and Materials and methods for details). First, we computed neural representational dissimilarity matrices (RDM) that reflected trial-by-trial variation in activation patterns throughout the scanning task. Second, we used post-scan ratings to construct behavioural RDMs that reflected, respectively, between-trial differences in perceived distress experienced by the co-participant, and between-trial differences in how threatened the participant felt themselves during the scan. Finally, in the third step, we estimated the second-order similarity between neural and behavioural RDMs. This similarity metric allowed us to assess, for each ROI, whether trial-by-trial multivoxel patterns during the scan primarily represented the co-participant’s distress, or the threat to oneself. Importantly, it allowed us to determine whether and to what degree neural representations of other’s distress and of threat to the self were associated with helping behaviour. Results show that, regardless of threat imminence, the similarity between neural and threat RDMs in the left amygdala (*β*=4.41, *se*=1.35, *t*=3.27, *p*=.006) and left insula (*β*=2.46, *se*=0.97, *t*=2.53, *p*=.047) was positively associated with helping behaviour (Figure 6; Table S3). In other words, the more strongly these brain regions, especially the amygdala, represented the threat to oneself, the more frequently the participant decided to help.

**Figure 5.**
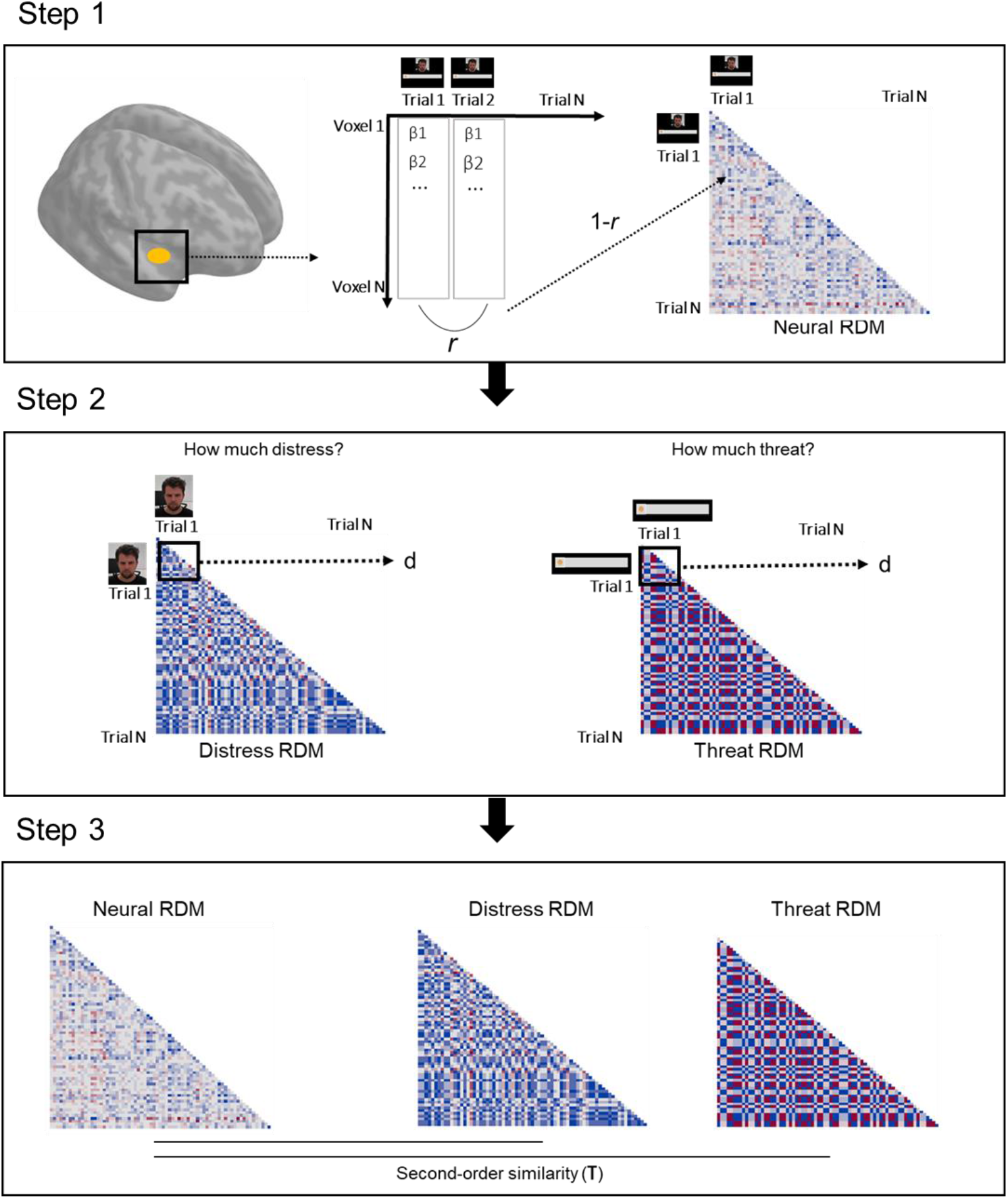
Schematic of the RSA pipeline. On step 1, we extracted the vector of trial-by-trial betas for each voxel in a given ROI. We then calculated the correlation (Pearson r) between all trial pairs. These correlation values were inverted (1-r) and used to create a trial by trial matrix, wherein each cell represents how correlated activation across all voxels of the ROI was in each trial pair (neural representational dissimilarity matrix, RDM). On step 2, post-scan ratings of the co-participant’s distress in each unique clip were used to construct a trial-by-trial matrix, wherein each cell contained the Euclidean distance between the rating of each pair of clips (distress RDM). A similar method was used with the ratings of threat to the participant (threat RDM). On step 3, the second-order similarity between the neural RDM and distress RDM, and between the neural RDM and threat RDM were calculated using a ranked correlation method (Kendall’s tau).

**Figure 6.**
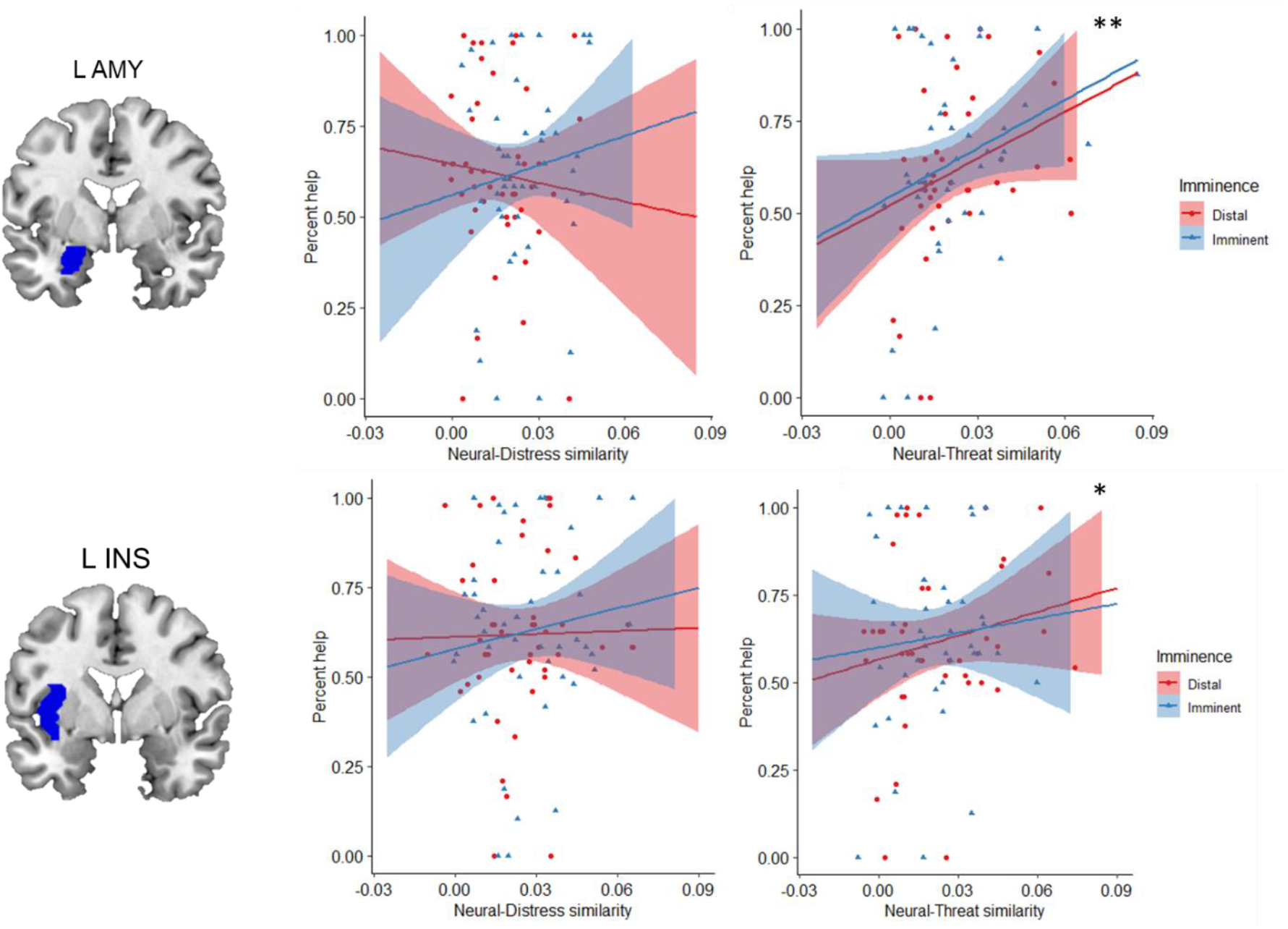
Regardless of threat imminence, the similarity between neural and threat RDMs in the left amygdala and insula predicted higher frequency of helping decisions. \**p* = .047; \*\**p* = .006

## Discussion

Our overarching goal with this study was to determine how one’s own defensive responses influence decisions to help others under threat. Our findings strongly suggested that neural circuits that coordinate fast avoidance responses from self-directed threats (reactive fear circuits) (*12*) also underlie the protection of others in dangerous situations. More, the extent to which key defensive regions (amygdala and insula) represent the threat to the self (and not others’ distress) predicts more frequent helping decisions.

### Reactive fear circuits promote self- and -other defence

Previous work has shown that increased imminence of an other-directed threat facilitates helping behaviour, suggesting that the same way threat imminence triggers active avoidance (*i.e*., fight or flight) from self-directed threats, it may promote defensive helping when others are under threat (*17*). Here, we examined the neural basis of this effect. We found that multivoxel activation within the defensive circuitry (*i.e.,* amygdala, ACC, insula, hippocampus and OFC) tracked with the level of threat. Also, we found dissociable local patterns of activation prior to helping as a function of threat imminence within the insula, hippocampus and dorsal ACC. These findings are consistent with previous work highlighting the role of the hippocampus and dACC as key regions optimizing escape decisions within cognitive and reactive fear circuits, respectively (*21*). Importantly, we found that average activation within several regions of the defensive circuitry, namely the in the PAG, insula, hippocampus, dACC, OFC and vmPFC, differed between trials leading up to helping or not helping decisions. Specifically, consistent with our predictions, activation in the insula, ACC and PAG was higher before decisions to help, whereas activation in the hippocampus, vmPFC, and OFC was higher prior to decisions not to help. Therefore, although overall we did not find higher frequency of helping decisions under imminent than distal threat, our results suggest that greater engagement of reactive fear circuits facilitated helping behaviour. These results are in line with prior findings suggesting that acute defensive states coordinated by reactive fear circuits promote helping under threat (*17*). Of note, the differential activation of cognitive and reactive fear circuits based on the subsequent decision (help versus not help) was only found when decisions were made under distal threat (that is, when responses were prompted in the beginning of the trial). Also, contrary to our predictions, when comparing the average activation during distal and imminent threats independently of the decision, we found that the bilateral insula and ACC were more active during distal relative to imminent threats, whereas the bilateral IFG and OFC were more active during imminent threats. These results are opposite to those from prior studies that manipulated the imminence of self-directed threats (*28, 30, 31, 33*). It is possible, however, that the disparity between our and previous findings is due to methodological reasons and does not reflect true differences in the processing of self-versus other-directed threats. In our paradigm, every trial started with a distal threat (the circle was static on the left) that always evolved into an imminent threat (the circle would move to the right, and the shock would be administered at the end of the trial). This may have made the threat overly predictable (especially given the high number of trials required for fMRI), engaging reactive fear circuits to a greater extent during the distal phase of the trial.

Nonetheless, our results obtained in a paradigm wherein the threat was directed at another person (the co-participant) were overall consistent with previous fMRI research investigating neural responses to self-directed threats as a function of imminence, suggesting a parallel between responses to self- and other-directed threats. This self-other parallel has been demonstrated in other threat-related processes, such as learning (*34*). Importantly, in line with previous demonstrations that acute defensive states may promote prosocial outcomes (*17*), we found that greater engagement of reactive fear circuits may facilitate helping of others in a threatening situation.

### The neural representation of threat to the self predicts helping

To decide whether to help another person in a dangerous situation, one must consider not only their distress, but also the threat in the environment. Here, we determined how the brain, in particular the defensive circuitry, represents another’s distress and threat when they are simultaneously present, and how those neural representations guide behaviour. To do so, we collected ratings of both the co-participant’s distress, and of how threatened the participant felt during the scan. Crucially, these ratings were obtained after the scan, allowing us to obtain a behavioural metric of how participants independently represented another person’s distress and the threat value of the situation. Performing these ratings after the scan also enabled us to avoid explicitly priming participants to consider those distress and threat cues during the scan, which could have influenced their behaviour and neural responses. Using behavioural representations of distress and threat, we assessed the extent to which each brain region in the defensive circuitry represented those cues, and its association with helping decisions. Our results showed that, regardless of threat imminence, the more the left amygdala and left insula represented the threat to oneself, the more participants decided to help.

The association between representation of threat and helping was particularly strong in the left amygdala. The amygdala has for long been known to have a pivotal role in the acquisition and expression of defensive responses in mammals (*35*). For instance, in both humans and rodents, it has been shown to coordinate switches between defensive states across through the communication between its basolateral nucleus (BLA) and oxytocin(OT)-sensitive neurons in the central amygdala (CeA) (*22*). Our present results suggest that the amygdala’s role in defensive responding may also be relevant for helping behaviour. This is consistent with previous work in animals. In rodents, CeA activation by OT not only enables the transition from freezing to fight-or-flight, but has also been shown to trigger offspring care behaviours in females (*10*), and to enhance maternal aggression (*9*). It has additionally been demonstrated that the administration of benzodiazepines, drugs with a known effect on CeA (*36*), impairs helping behaviour in rats (*8*). Taken together with these reports, our findings in humans suggest that amygdala-mediated defensive processes may also enable the provision of care to others, here in the form of helping. Of note, our effects were restricted to the left amygdala, which is consistent with several others demonstrations of hemispheric specialization in amygdala function in emotional and pain processing (*37*–*41*).

We found no evidence that the representation of other’s distress in any ROI was associated with helping, including in brain regions that have previously linked with empathy for pain and distress states (ACC and insula)(*42, 43*). Relatedly, our behavioural analysis revealed that, contrary to previous research (*44, 45*), empathic concern was negatively associated with the percentage of helping decisions. These findings are at odds with prior accounts of altruistic responding, which suggest empathy for distress is the key proximal mechanism driving helping behaviour (*6, 7*). Several explanations may account for lack of evidence of an association between neural representation of distress and helping in our data. One is that the behavioural ratings, which were collected after the scan, were not sensitive enough to reflect variations in perceived distress between clips of the confederate. However, this seems unlikely, given that we found significant effects of imminence and threat level on ratings of distress. Another possibility is that distress is mainly represented in brain regions not included in our analysis. The amygdala, insula and ACC would be prime candidate regions to represent another individual’s distress, in light of previous research on empathy (*46*–*49*), but we did not detect an association between the degree to which these regions represented distress and helping behaviour. Other potential regions would be those previously implicated in mentalizing, such as the temporo-parietal junction (TPJ) (*47, 50, 51*). At an exploratory level, we repeated the RSA analysis within anatomical masks of the left and right TPJ, but did not find evidence that the representation of distress in these regions were related to helping behaviour (see SI). Additional research at the whole brain level is necessary to further assess the representation of distress in the brain, and its impact on helping behaviour under threat. In any case, our present results suggest that, even if perceiving distress/need in others to some extent triggers altruistic motivation (*3*), the ability to provide help may ultimately rely on the activation of circuitry implicated in self-defence.

In summary, our results point to a parallel between responses to self- and other-directed threats, and suggest that the engagement of reactive fear circuits facilitates helping of others. Importantly, we showed that the extent to which the amygdala represents the threat to self (and not other’s distress) predicts helping decisions. These results challenge the idea that empathy for distress is the only proximal mechanism motivating helping decisions, and that overriding self-defensive responses is necessary to help others under threat. Rather, in dangerous situations, one’s own response to the threatening event may enable defensive helping of others, possibly through the activation of neural mechanisms subserving both individual defense and offspring care in mammals.

## Materials and Methods

This study has been pre-registered (https://osf.io/yvufn) and any deviations from the pre-registration are justified in the Supplementary information (SI). Data and code will be made available at the OSF project page (osf.io/9cuva).

### Participants

Forty-nine healthy volunteers (M=24.29, SD=4.78) participated in the experiment. Participants were recruited via flyers posted on- and off-campus, and local online recruitment systems. All participants were right-handed, had normal or corrected-to-normal vision, and were screened for history of psychiatric or neurological diagnoses, current medication, brain injuries and substance abuse. Participants provided informed consent prior to the experimental, and were compensated for their participation. The study was approved by the regional ethics board in Stockholm, Sweden.

### fMRI helping under threat task

In each testing session, a participant and a confederate (henceforth, co-participant) were informed the experiment comprised two parts (only one of those parts involved an MRI scan), which would be randomly assigned to each one by flipping a coin. Participant and co-participant were then accompanied to separate testing rooms (the actual participant was taken to the MRI area) and did not interact again during the experiment (details about testing procedures, post-task questionnaires and debriefing are available in the SI).

In the MRI, participants performed a task modified from previous work (*17*) wherein they made trial-by-trial decisions about whether or not to help the co-participant avoid aversive electrical shocks to the wrist, at the risk of also being shocked (Figure 1A). Threat imminence was manipulated by varying the spatial position and movement of a visual cue signaling varying levels of threat on a computer screen. Respectively, a green circle signaled no threat (no shocks), a yellow circle signaled moderate threat (1 upcoming shock) and the red circle signaled high threat (2 upcoming shocks). In addition, a webcam feed of the co-participant was presented on the screen throughout the task. Unbeknownst to the participant, the video feed was in fact pre-recorded, and edited to select unique clips for each trial of the task.

Participants were informed that, throughout the experiment, they and the co-participant would see the same screen. Each trial started with a static cue on the left side of the screen (4 s), which then moved to the right (4 s). In shock trials, the co-participant would be administered an aversive shock to the wrist when the cue reached the right end of the screen, unless participants decided to help him. To decide whether they wanted to help the co-participant avoid the upcoming shock, participants made forced-choice responses by pressing 1 (Help) or 2 (Don’t help) on an MRI-compatible button box as soon as the response slide was displayed (1.25 – 1.75 s). Responses were prompted sometimes in the beginning of the trial, when the visual cue was static on the left side (distal threat), and other times at the end of the trial, after the visual cue had moved to an endpoint on the right, and thus immediately before shock delivery (imminent threat). Outcomes of participants’ decisions were as follows: if they chose not to help, the co-participant would always receive a shock; if they chose to help, there would be around 70% probability of both participant and co-participant receiving 1 shock (in moderate threat trials) or 2 shocks (high threat trials). Shocks were administered on the left ankle. Participants were instructed they should respond as quickly as possible. Also, to discourage missed responses, they were informed that a shock would be delivered to both participants (with 100 % chance) whenever a response was not detected. Finally, to balance the number of helping and non-helping trials during the scanning session, participants were informed that they would have a pre-set number of times they could help on each run, and thus they should try to balance, per run, the number of times they helped and not helped. In reality, participants could help on as many trials as they wished. Shock administration always happened at the end of the trial, and participants were able to see the outcome of their decisions on the screen (i.e., the co-participant receiving or not receiving a shock; 4 s).

Safe trials followed an identical structure, with response slides presented at distal or imminent stages in relation to the end of the trial. However, participants were instructed that no shocks would be given and they should arbitrarily choose to press 1 or 2 when the response slide was displayed. It was made clear to them that their choice would have no consequences for them or the co-participant.

The task included 144 trials split into 8 functional runs (approx. 8 minutes). Each run comprised 18 trials, 9 distal and 9 imminent, and 6 of each threat level (resulting in 24 trials per condition, in total). Distal and imminent trials were presented in blocks, and the order of blocks was in each run. Within each block, safe, moderate threat and high threat trials were randomized. The order of functional runs was randomized across participants. The task was programmed and delivered using E-prime 3.0 (Psychology Software Tools, Inc., www.pstnet.com).

### Ratings

After the scan, participants were taken to a different testing room and asked to complete a follow-up task. Here, all video clips showed during the scanning task were presented to participants, in random order. Participants were informed that these had been recorded during the scan, and that their task now was to, for each clip, rate the level of distress, anxiety or concern they perceived in the co-participant, on a 9-point scale. Participants were also presented images of the visual cues at distal (left side of the screen) and imminent positions (right side of the screen), and asked to rate on a 9-point scale how threatened they felt during the scan, whenever they saw those images (Figure 1B). Ratings of distress and threat were presented in separate blocks, and the order was randomized.

### fMRI acquisition and preprocessing

Participants were scanned in a single session at the Stockholm University Brain Imaging Center (SUBIC), using a 3T Siemens scanner with a 64-channel head coil. First, a high resolution t1-weighted anatomical scan was obtained (TR = 2300 ms, TE = 2.98ms; FoV = 256mm, flip angle = 9°, 192 axial slices of 1 mm isovoxels), followed by 8 functional runs, of about 8 min each. Functional images were acquired with an echo-planar T2*-weighted imaging sequence with whole-brain coverage while participants performed the fMRI task (TR = 1920ms, TE = 30ms, FoV = 192 mm, flip angle = 70°, 62 interleaved slices of 2mm isovoxels, acceleration factor of 2).

Preprocessing of fMRI data was done using SPM12 (Wellcome Trust Centre for Neuroimaging, www.fil.ion.ucl.ac.uk), and included slice timing correction, realignment to the volume acquired immediately before the anatomical scan (i.e., the first image of the first functional sequence) using six parameter rigid-body transformations, coregistration with the structural data, normalization to standard space using the Montreal Neurological Institute (MNI) template with a voxel size of 2 x 2 × 2 mm, and smoothing using a Gaussian kernel with an isotropic full-width-half-maximum of 4 mm (*52, 53*). Finally, a high-pass filter with a cutoff of 128 s was applied to remove slow signal drifts.

### Statistical analysis

#### Behavioural data

Behavioural data was in general analyzed using Generalized Linear Mixed Models (GLMMs), an approach that accounts for variation in the dependent variable that is explained by random sampling of, for instance, participant or trial number (random effects), in addition to the independent variables (fixed effects). Mixed-effects approaches have further been proposed to increase the generalizability of research findings to other individuals and stimuli (Yarkoni, 2020).

Our main behavioural variable was helping behaviour, which was operationalized as the percentage of helping responses throughout the task. We modelled helping percentage using GLMMs as a function of threat imminence, threat level, and imminence by threat interaction (fixed effects). The subject was added as a random effect, with random intercept and slope per threat imminence and level. In a separate model, we also added a threat imminence*level*empathic concern interaction, following previous indications that threat imminence may affect helping behaviour more strongly in individuals with higher caregiving tendencies. To account for the possibility that behaviour varied throughout the experiment, we also performed a mixed effects logistic regression on single trial dichotomous responses (help or no help), including the trial number as a fixed effect in addition to the other fixed effects. Finally, following recent recommendations to consider within-individual effect sizes (*54*), we also calculated the difference between number of helping decisions under imminence and distal threat, per individual.

Reaction times were averaged per condition, and analysed in a GLMM with threat imminence, threat level, and imminence by level interaction as fixed effects, and the subject as a random effect (intercept and slope). Similarly, post-task ratings of other’s distress and threat to self were analysed in a GLMM with threat imminence, threat level, and imminence by level interaction as fixed effects, and the subject as a random effect (intercept and slope).

#### Imaging data

##### a. First-level analysis

First-level analysis was performed in SPM12 and was based on the general linear model. Time-series of each voxel were normalized by dividing the signal intensity of a given voxel at each point by the mean signal intensity of that voxel for each run and multiplying it by 100. Resulting regression coefficients thus represent a percent signal change from the mean. Regressors were created by convolving the train of stimulus events with a canonical hemodynamic response function. Three different GLMs were estimated based on the goal of the analysis. For assessing differences based on threat imminence and level, 6 regressors of interest were modelled corresponding to the time window of the visual threat cue (distal safe, distal 1 shock, distal 2 shocks, imminent safe, imminent 1 shock and imminent 2 shocks). These regressors were defined based on the position of the threat cue on the screen (static = distal; approaching = imminent), and not relative to when the participant made a decision. In addition, 8 regressors of no interest were added in the model, corresponding to the time window of the response and the outcome, plus the 6 motion parameters estimated during realignment.

To assess differences in neural response based on the type of decision, another model was estimated with 6 regressors of interest: help distal 1 shock, help imminent 1 shock, help distal 2 shocks, help imminent 2 shocks, no help distal, no help imminent. Here, distal and imminent refer specifically to when the decision was prompted in the trial. Of note, due to the reduced number of no helping trials for some participants, the “no help” regressor included both not help decisions, and decisions made during safe trials, wherein no shocks were given. In addition, 8 regressors of no interest (response, outcome and 6 motion parameters) were added.

A third model was created to enable subsequent trial-by-trial Representational Similarity Analysis (RSA), wherein one regressor was estimated for each individual trial, modelling the time window of the threat cue.

##### b. Regions of interest (ROIs)

Given our focus on defensive brain circuitry, our analysis targeted pre-specified ROIs that were anatomically defined, including the left and right amygdala, left and right hippocampus, left and right insula, midbrain, left and right ACC, left and right vmPFC, left and right vlPFC (Figure 3). ROIs were defined on the Wake Forest University (WFU) Pickatlas toolbox (http://www.fmri.wfubmc.edu/cms/software) (Maldjian et al., 2003).

##### c. MVPA and SVR

Beta values derived from first-level analyses were used in multivariate analyses, including multivoxel pattern analysis (MVPA) searchlight, support vector regression (SVR), and representational similarity analysis (RSA). Spatially distributed patterns of activation across voxels can reveal distinguishable neural responses between experimental conditions even in the absence of significant average activation differences in single voxels, making multivariate approaches more sensitive than conventional univariate analysis (*55*).

To identify local activation patterns that distinguish between distal and imminent threats, we used an MVPA Searchlight implemented in The Decoding Toolbox (TDT). A spherical searchlight (radius 12 mm) was moved throughout each participant’s data and, at each searchlight center, a support vector machine algorithm was trained to discriminate activation patterns in response to distal and imminent threats. Training was done iteratively on each 7 functional runs and tested on the 8th (leave-one-out cross-validation). Resulting percentage score at each voxel for participant was calculated and display in individual accuracy maps. Accuracy maps were then analysed at the group level in a one-sample T test implemented in SPM12. A similar approach was taken to the identification of local activation patterns that discriminated helping decisions during distal and imminent threats. Additionally, a SVR was used to identify local patterns that showed a continuous linear association with threat level (safe, 1 shock, 2 shocks). As for the classification searchlight, individual accuracy maps were analysed at the group level in a one sample t test. Group results were thresholded at voxelwise FWE < .05 and only clusters with more than 10 voxels were further considered.

##### d. Univariate analysis

We also performed GLM-based univariate analyses. We analysed BOLD signal to the threat cues, regardless of decision, in a threat imminence (distal, imminent) by threat level (safe, 1 shock, 2 shocks) repeated-measures ANOVA. We also analysed BOLD signal to threat cues prior to the decision in a decision type (help, not help) by threat imminence (distal, imminent) ANOVA. Full results for these analyses are available in supplementary material. Univariate analysis results were first thresholded at p<.001 uncorrected. With this threshold, clusters with more than 10 voxels were significant with FWE-corrected p <.05.

##### e. RSA

One of the goals of the study was to characterize representations of other’s distress and of threat to oneself by defensive regions, and determine its relation to helping behaviour. RSA was done separately for imminent and distal trials, and comprised 3 steps (Figure 5). On the first step, we modelled each trial in the first-level analysis (in SPM12), in order to estimate one beta coefficient per trial. Then, for each ROI, we extracted betas from each voxel in each trial to estimate the correlation of beta-values (expressed in r values) between all voxels per trial. Resulting r values were used to construct a representational similarity matrix across all trials, that reflects the correlation between all voxels in each trial. This matrix was then transformed (1-r) to reflect dissimilarity instead of similarity (RDM). On the second step, distress ratings provided after the scan on the unique video clips shown in each trial were used to construct a dissimilarity matrix that reflects the difference in perceived distress of the co-participant between trials (expressed in Euclidean distances). Post-scan threat level ratings were used in an identical manner to construct a dissimilarity matrix that reflects between-trial differences in how threatened the participant felt during the scan. Finally, on the third step, we estimated the second-order similarity (kendall’s τ) between neural and behavioural RDMs. In a nutshell, this similarity metric allowed us to assess, for each ROI, whether trial-by-trial multivoxel patterns during the scan primarily represented the co-participant’s distress, or the threat to oneself. Importantly, it allowed us to determine whether neural representations of other’s distress and of threat to oneself were associated with helping behaviour. To do so, second-order similarity values were entered in a linear model predicting average helping percentage during the scan. Predictors in this model were the similarity between neural and threat RDM, the similarity between neural and distress RDM, as well as threat imminence. Fourteen linear models were estimated, one for each ROI (i.e., left and right amygdala, left and right hippocampus, left and right insula, midbrain, left and right ACC, left and right vmPFC, left and right vlPFC). False discovery rate (FDR) correction was applied to adjust the p value of all coefficient estimates, across all 14 models. FDR-corrected p values below α=.05 were considered significant. Beta value extraction was performed in Matlab, and all remaining steps and analyses of the RSA were performed through custom-made scripts in R.

## Acknowledgments

We thank Dean Mobbs, India Morrison, Björn Lindström, Artin Arshamian, and Henrik Ehrsson for their helpful and insightful comments on our manuscript.

## Author contributions

J.B.V. designed and performed the research, and analyzed the data; J.B.V. and A.O. wrote the paper.

## Data and materials availability

Anonymized data, code, and materials used in the study will be available on the OSF project page (osf.io/9cuva).

