## Supplements for "Neural defensive circuits underlie helping under threat in humans"

### Supplemental Information

1. Experimental procedures

The experiment involved one session. Upon arrival, both participant and co-participant (i.e., confederate) were welcomed into the lab and told that the experiment comprised two parts and each one of them would only perform one. The parts were assigned through a fake coin flip to give the impression of randomization. Participant and co-participant were then accompanied to separate rooms to receive more detailed instructions.

Prior to entering the MRI, written informed consent was obtained, and electrodes for administration of electrical shocks were placed on participants' left ankle. Participants were then given a short practice block on a computer, to allow them to familiarize with the timing of responses. After the practice, participants were taken into the MRI scanner room.

Once in the scanner, shock intensity was individually calibrated using a standard work-up procedure. Participants were asked to select an intensity level that was "not painful, but very uncomfortable" and that it should be something that "if they could avoid, they would rather avoid". It was emphasized to participants that, during the experiment, they would be able to avoid the shock if they so desired.

Participants were then given written instructions for the first block of the task. They were informed they would see the co-participant via webcam and they would both be presented the same stimuli on the screen. The first block consisted of 3 trials (safe, 1 shock and 2 shocks), wherein the participant was simply required to pay attention to the screen; they would not be asked to make any responses and would not be given any shocks. The goal of this block was to allow participants to see the consequences of safe and threat trials to the co-participant. This was meant to discourage them from making "test responses" in the first trials of the actual task (for instance, decide not to help just to see whether the co-participant would indeed receive a shock).

Thereafter, participants were given written instructions for the rest of the task. They were explained that, on each trial, they would be asked whether they wanted to help the co-participant avoid the shock(s) or not, and what would be the outcomes of their decisions. Participants were also informed that: 1. the co-participant was not aware that shock administration was decided by them, 2. they should try to balance out the number of help and not help decisions, since there was a preset number of times they could help in each run, 3. they would not swap places with the co-participant afterwards, 4. they would not meet the co-participant again after finishing the experiment, and 5. their behavior during the task would not be observed nor filmed.

After the scan, participants were taken to another testing room to perform the ratings task. At the end, they were asked to fill out questionnaires, including post-tasks questions designed to assess whether the instructions and cover-story were believable (e.g., To what extent do you think the shocks were controlled by you? How authentic did the co-participant seem to you?). Lastly, they were fully debriefed.

*1.1. Post-task and questionnaire measures*

After the fMRI helping under threat task and the ratings task, participants completed a series of questionnaires to assess individual differences in empathy and threat sensitivity, including the Interpersonal Reactivity Inventory (Davis, 1983), the Trait Fear Questionnaire (Kramer et al., 2020), the, and the Triarchic Psychopathy Measure (Patrick et al., 2009). Additionally, participants completed a series of post-task questions designed to assess the believability of the experiment.

2. Univariate analysis

*2.1. Table S1. Threat imminence X Threat level Anova*

| **Main effect of threat imminence** | | | | | |
|---|---|---|---|---|---|
| | **R/L** | **k** | **x, y, z** | **F** | **BA** |
| Insula | R | 36 | 42, 2, -6 | 12.76 | 13 |
| Insula | L | 113 | -28, 12, -20 | 16.29 | 47 |
| IFG | R | 1410 | 50, 20, -10 | 25.43 | 47 |
| vmPFC | L | 334 | -8, 32, -16 | 17.43 | 11, 25, 10 |
| OFC | L | 37 | -18, 56, -8 | 12.62 | 11 |
| **Main effect of threat level** | | | | | |
| | **R/L** | **k** | **x, y, z** | **F** | **BA** |
| IFG | R | 72 | 36, 24, 8 | 20.25 | 13, 45, 47, 44, |
| vmPFC | L | 443 | 0, 34, -16 | 27.35 | 10, 11, 32, 25, 9 |
| **Threat imminence*level** | | | | | |
| | **R/L** | **k** | **x, y, z** | **F** | **BA** |
| Insula | R | 768 | 38, 20, 6 | 28.14 | 13 |
| Insula | L | 521 | -30, 24, -4 | 20.58 | 13 |
| IFG, OFC | L | 593 | -40, 32, -16 | 23.72 | 11 |
| OFC | R | 407 | 8, 32, -20 | 23.43 | 11 |
| ACC | R | 278 | 4, 32, 20 | 18.29 | 24 |
| OFC, IFG | R | 52 | 34, 40, -12 | 13.18 | 11, 47 |

*2.2. Table S2. Type of decision X Threat imminence Anova*

| **Main effect of type of decision** | | | | | |
|---|---|---|---|---|---|
| | **R/L** | **k** | **x, y, z** | **F** | **BA** |
| Hippocampus | L | 128 | -32, -18, -14 | 17.85 | |
| Hippocampus | R | 58 | 28, -16, -16 | 16.01 | |
| Insula | L | 638 | -40, 8, 6 | 24.10 | 13, 47, 44, 45, 22 |
| Insula | R | 949 | 46, 14, -2 | 30.43 | 13, 47, 44, 45, 22 |

| | | | | | |
|---|---|---|---|---|---|
| ACC | R | 218 | 2, 22, 28 | 24.33 | 32, 6, 24, 8 |
| IFG | L | 104 | -38, 28, -14 | 12.38 | 21, 38, 47, 22 |
| vmPFC | L | 768 | 0, 44, -18 | 24.74 | 11, 25, 10, 32 |
| OFC | R | 129 | 36, 54, -14 | 14.50 | 11, 10 |
| **Main effect of threat imminence** | | | | | |
| | **R/L** | **k** | **x, y, z** | **F** | **BA** |
| Insula | R | 40 | 36, -20, 8 | 21.07 | 13 |
| | | 56 | 38, 8, 4 | 17.93 | 13 |
| | | 45 | 32, 20, 14 | 17.09 | 13, 45 |
| IFG | L | 211 | -50, 42, -8 | 29.37 | 47, 10 |
| vmPFC | L | 447 | -2, 54, -14 | 36.39 | 11 |
| **Type of decision*threat imminence** | | | | | |
| | **R/L** | **k** | **x, y, z** | **F** | **BA** |
| Midbrain | | 60 | 2, -32, -4 | 16.85 | |
| Insula | R | 192 | 38, -16, -2 | 20.62 | 13, 47, 22, 44, 6, 45, 21 |
| Hippocampus | R | 38 | 30, -14, -20 | 15.03 | |
| Insula | L | 595 | -36, -8, -4 | 18.68 | 13, 22, 44, 6, 47, 45 |
| Dorsal anterior cingulate | L | 151 | -2, 14, 30 | 16.13 | 24, 6, 32, 5, 4 |
| Insula | R | 748 | 30, 20, -8 | 18.18 | 47, 13, 22, 44, 6, 45, 21 |
| IFG/orbital gyrus | L | 46 | -34, 36, -10 | 11.47 | 11 |
| Ventral med frontal gyrus | L | 238 | 0, 46, -20 | 19.94 | 11, 10, 25 |

3. RSA

Table S3. Estimates from models for each ROI, predicting helping percentage throughout the scan as a function of neural-distress similarity, neural-threat similarity, and threat imminence (all *p* values across all models were FDR-corrected).

| | **Estimate** | **Std. Error** | **t value** | **p (FDR-corrected)** |
|---|---|---|---|---|
| **L amygdala** | | | | |
| Intercept | 0.536 | 0.054 | 9.823 | <.00001 |
| Threat | 4.407 | 1.348 | 3.269 | **.006*** |
| Distress | -1.292 | 2.049 | -0.630 | .887 |
| Imminence | 0.040 | 0.055 | 0.731 | .837 |
| **L insula** | | | | |
| Intercept | 0.531 | 0.059 | 8.883 | <.00001 |
| Threat | 2.461 | 0.974 | 2.525 | **.047*** |
| Distress | 1.286 | 1.552 | 0.828 | .837 |
| Imminence | 0.0129 | 0.052 | 0.246 | .887 |
| **L ACC** | | | | |
| Intercept | 0.649 | 0.057 | 11.257 | <.00001 |
| Threat | 0.216 | 0.940 | 0.230 | .887 |
| Distress | -1.339 | 1.304 | -1.026 | .742 |
| Imminence | 0.018 | 0.054 | 0.348 | .887 |
| **L hippocampus** | | | | |
| Intercept | 0.620 | 0.067 | 9.233 | <.00001 |
| Threat | 0.042 | 1.564 | 0.027 | .978 |
| Distress | 0.104 | 1.642 | 0.063 | .968 |
| Imminence | 0.017 | 0.054 | 0.324 | .887 |
| **Midbrain** | | | | |
| Intercept | 0.722 | 0.066 | 10.936 | <.00001 |
| Threat | -2.196 | 1.694 | -1.296 | .573 |
| Distress | -2.622 | 1.753 | -1.495 | .451 |

| | | | | |
|---|---|---|---|---|
| Imminence | 0.040 | 0.055 | 0.736 | .837 |
| **L vmPFC** | | | | |
| Intercept | 0.589 | 0.059 | 9.842 | <.00001 |
| Threat | 0.752 | 1.010 | 0.744 | .837 |
| Distress | 0.627 | 1.495 | 0.419 | .887 |
| Imminence | 0.017 | 0.054 | 0.316 | .887 |
| **L vlPFC** | | | | |
| Intercept | 0.589 | 0.059 | 9.842 | <.00001 |
| Threat | 0.752 | 1.010 | 0.744 | .837 |
| Distress | 0.627 | 1.495 | 0.419 | .887 |
| Imminence | 0.017 | 0.054 | 0.316 | .887 |
| **R amygdala** | | | | |
| Intercept | 0.695 | 0.062 | 11.170 | <.00001 |
| Threat | -1.651 | 1.584 | -1.042 | .742 |
| Distress | -1.319 | 1.411 | -0.935 | .797 |
| Imminence | 0.017 | 0.053 | 0.332 | .887 |
| **R insula** | | | | |
| Intercept | 0.603 | 0.060 | 10.015 | <.00001 |
| Threat | 0.436 | 0.976 | 0.447 | .887 |
| Distress | 0.189 | 1.676 | 0.113 | .946 |
| Imminence | 0.020 | 0.054 | 0.376 | .887 |
| **R ACC** | | | | |
| Intercept | 0.541 | 0.060 | 8.883 | <.00001 |
| Threat | 1.781 | 1.279 | 1.392 | .512 |
| Distress | 1.617 | 1.456 | 1.110 | .739 |
| Imminence | 0.014 | 0.053 | 0.276 | .887 |
| **R hippocampus** | | | | |
| Intercept | 0.677 | 0.063 | 10.677 | <.00001 |
| Threat | -1.451 | 1.432 | -1.012 | .742 |

| | | | | |
|---|---|---|---|---|
| Distress | -0.704 | 1.350 | -0.521 | .887 |
| Imminence | 0.017 | 0.054 | 0.326 | .887 |
| **R vmPFC** | | | | |
| Intercept | 0.612 | 0.058 | 10.490 | <.00001 |
| Threat | 0.933 | 1.055 | 0.884 | .820 |
| Distress | -0.467 | 1.345 | -0.347 | .887 |
| Imminence | 0.012 | 0.047 | 0.272 | .887 |
| **R vlPFC** | | | | |
| Intercept | 0.603 | 0.066 | 9.134 | <.00001 |
| Threat | -0.242 | 1.405 | -0.172 | .916 |
| Distress | 1.118 | 1.670 | 0.669 | .876 |
| Imminence | 0.0207 | 0.054 | 0.381 | .887 |

Table S4. Exploratory examination of association between neural-distress and neural-threat similarity, and helping in the temporo-parietal junction (TPJ) (anatomically defined ROI based on aal, including supramarginal and angula gyri). Estimates from linear models for left and right TPJ.

| | Estimate | Std. Error | t value | p |
|---|---|---|---|---|
| **L TPJ** | | | | |
| Intercept | 0.552 | 0.051 | 10.756 | <.00001 |
| Threat | 1.669 | 0.967 | 1.725 | 0.088 |
| Distress | 1.103 | 1.492 | 0.739 | 0.462 |
| Imminence | 0.014 | 0.053 | 0.269 | 0.789 |
| **R TPJ** | | | | |
| Intercept | 0.559 | 0.057 | 9.859 | <.00001 |
| Threat | 1.407 | 0.842 | 1.671 | 0.098 |
| Distress | 0.585 | 1.491 | 0.392 | 0.696 |
| Imminence | 0.018 | 0.054 | 0.339 | 0.735 |

4. Deviations from pre-registration

A pre-registration for this study can be found at https://osf.io/yvufn. Some changes were implemented in relation to the pre-registration, namely:

1. Correction for multiple comparisons in the analysis of fMRI data was done via Family-Wise Error (FWE) and not False-Discovery Rate (FDR). FWE was deemed a more conservative approach, and adequate to our focus on regions-of-interest.
2. Group results for MVPA searchlight analyses were done with t-test and not permutations. This alteration was in line with recommended procedures (Hebart et al., 2015). Median maps were assessed to confirm they were in agreement with the parametric results.
3. Parametric modulator analysis was performed (results available on https://osf.io/9cuva/) but not included in the manuscript. The reason for this was that several subjects did not have enough responses on some conditions for the analysis to be reliable, resulting in only 28 subjects being included. Instead, we estimated another GLM, in which all 'not help' and 'safe' decisions were considered not help and contrasted with help decisions in a group-level type of decision X threat imminence anova (reported above and in the manuscript).
4. Out of 49 participants, only one participant stated categorically not believing in the cover story. We deemed it to be a more conservative approach to keep this participant in the analysis, especially since it was unclear at what point of the experiment they started to have doubts.
5. We pre-registered an exploratory whole-brain RSA Searchlight analysis that is still ongoing, and is thus not included in the present manuscript.
